# Unveiling mode of action of anthelmintics in *Caenorhabditis elegans* with SydLab™, an on-a-chip automated and high-content screening system

**DOI:** 10.1101/2025.05.15.654210

**Authors:** Thomas B. Duguet, Coralie Belgrano, Morgane Bourgeois, Alexandre Vernudachi, Laurent Mouchiroud, Lucien Rufener

## Abstract

Anthelmintic resistance in parasitic nematodes presents a growing challenge to animal and human health, driving the need for innovative tools to accelerate drug discovery and mechanistic research. Here, we introduce SydLab™, a microfluidic-based, automated phenotypic screening platform that combines continuous imaging, machine vision, and computational analysis for high-content assessment of *Caenorhabditis elegans* responses to anthelmintics.

We systematically evaluated eight anthelmintic compounds spanning major chemical classes— albendazole, ivermectin, milbemycin oxime, emodepside, levamisole, tribendimidine, monepantel, and closantel—across multiple doses. Using wild-type (N2 Bristol) and mitochondrial stress-sensitive (*hsp-6::gfp*) strains, SydLab™ captured real-time, dose-dependent phenotypic profiles over 120 hours, measuring developmental growth, reproduction, motility, and morphology. Emodepside, monepantel, and macrocyclic lactones induced severe larval arrest, reduced worm volume, and distinct morphological changes consistent with neuromuscular paralysis. In contrast, albendazole and closantel showed limited effects at tested concentrations.

Machine learning-based shape classification revealed drug-specific morphological signatures, including coiling and cuticular damage, offering insights into compound modes of action. Validation with larval development and migration assays confirmed SydLab™’s high sensitivity and reproducibility in detecting subtle phenotypic and strain-specific responses.

Our study demonstrates how integrating microfluidics, automated imaging, and computational phenotyping enables precise dissection of complex drug-induced effects in nematodes. SydLab™ provides a scalable, high-throughput platform for anthelmintic screening and mechanistic studies, offering new avenues for antiparasitic drug discovery and resistance research.

**Author summary:** Resistance to anthelmintic drugs is a global concern threatening the control of parasitic nematodes that impact human and animal health. New tools are needed to better understand drug effects and accelerate the discovery of effective treatments. In this study, we present SydLab™, an automated microfluidic platform that combines real-time imaging and computational analysis to assess the impact of anthelmintic drugs on the model nematode *Caenorhabditis elegans*.

By monitoring development, reproduction, motility, and morphology, we identified unique drug-induced phenotypes linked to specific modes of action. Compounds like emodepside, ivermectin, and monepantel caused larval arrest and neuromuscular paralysis, while albendazole and closantel showed limited activity. Machine learning algorithms detected drug-specific morphological patterns, offering deeper mechanistic insights.

Our work highlights SydLab™ as a powerful high-throughput phenotyping system that can reveal subtle biological responses and support the discovery of next-generation anthelmintics. This approach strengthens our ability to investigate drug resistance and improve therapeutic strategies against parasitic nematodes.

## Introduction

Anthelmintics are a class of molecules whose use represents one of the primary strategies for combating parasitic diseases caused by helminthic species (nematodes, trematodes, and cestodes) in humans, animals, and plants. These molecules can either kill the target parasitic species or neutralize their natural behavior within their hosts, thereby indirectly leading to their expulsion (1,2).

One of the current research strategies for anthelmintics has focused on elucidating the modes of action of known molecules to define their targets and create new active derivative molecules (3–6). Additionally, considerable efforts have been directed towards the discovery of new active molecules through screening campaigns that employ various approaches, such as searching for natural products (7,8). This strategy has led to the development of promising compounds such as emodepside (9) and paraherquamides (10), or synthetic compounds from amino-aceto nitrile derivatives from which monepantel was introduced (11,12). Lastly, in the context of anthelmintic resistance reported for the majority of existing active families with a special concern for drugs used against parasites of veterinary importance (13,14), understanding these mechanisms is crucial for adjusting treatment methods or enhancing the search for active agents with innovative modes of action. However, the development of new anthelmintics is a lengthy and costly process that necessitates a strategic approach to achieve rapid and high-value results (15,16).

Among these approaches is the application of the latest technological advancements to parasites such as nematodes. In recent years, numerous innovations have enhanced screening processes by increasing throughput, automating reading and analysis processes (17,18), and integrating novel parameters such as the migration capacity of nematodes under treatment (19–21).

The model nematode *Caenorhabditis elegans* remains central to anthelmintic research (22) due to its ease of maintenance, well-characterized biology, and phylogenetic proximity to parasitic species—facilitating the discovery and mode-of-action studies of many drugs including commonly used anthelmintics (23–25). This *C. elegans*-based strategy continues to offer promising discoveries for the future, and it is in this context that SydLab™ technology developed by Nagi Bioscience SA has been integrated into the present study.

Most anthelmintics possess neuro-modulatory properties, particularly affecting neuromuscular communication through direct or indirect interactions with ion channels on nerves and muscles, leading to the inability of exposed worms to move (25–27). Motility analysis remains the principal tool for observing the effects of these compound families (28,29); however, in recent years, there has been significant progress in new methods for evaluating anthelmintic capacity *in vitro* based on robust phenotypic readings combining robotics, artificial intelligence, and microfluidics (5,30–34). Moreover, with the support of microfluidics, certain technical barriers are gradually being overcome, allowing for greater flexibility and the exploration of increasingly complex biological questions (35,36).

SydLab™ is a microfluidic-based robotic platform capable of performing automated high-content phenotypic analyses of *C. elegans*. It can conduct various assays, including the evaluation of developmental processes, reproductive mechanisms, morphogenesis, and chemically induced stress responses. This "nematode-on-chip" approach is designed to continuously evaluate these mechanisms using machine vision and machine learning, aspects of which could represent a major asset in research on any anthelmintic compound, regardless of its mode of action.

In this study, eight known anthelmintics (albendazole, ivermectin, milbemycin oxime, emodepside, levamisole, tribendimidine, monepantel, and closantel) were selected for their use in animal health (37,38), documented modes of action (39,40), and/or association with resistance phenomena (1,41). Two *C. elegans* strains, the wild-type N2 Bristol and the *hsp-6* transgenic strain, served as study models, with the latter enabling the investigation of the effects of the compounds on mitochondrial stress response in worms (42–44). These strains were incubated and exposed to increasing doses of each anthelmintic over time, from L1 larval stage to adult, allowing the collection of images and recording of each measured parameter throughout the period. Additionally, a recently developed migration test, also known as the "migration trap assay" (21,45,46), provides comparative elements with data obtained from SydLab™.

The entirety of these data can be viewed as an important starting point in characterizing active molecules, highlighting the complexity of the induced effects on nematodes, while proposing an innovative "nematode-on-chip" approach that could potentially be adapted to parasitic species.

## Results

### Dose-dependent negative effects of most of the tested anthelmintics on the morphological development of *C. elegans*

To establish a detailed characterization of the phenotypes induced by anthelmintic treatments in *C. elegans*, the SydLab™ system was fully leveraged to study their impact on developmental processes. Eight molecules from seven classes of anthelmintics were selected (Table 1) for their relevance to animal health, either due to their widespread use or well-defined modes of action. Throughout the study, the *hsp-6::gfp*/SJ4100 transgenic strain of *C. elegans* was employed, as it allows mitochondrial stress response levels to be correlated with fluorescence emission, which can be directly measured using SydLab™. Three replicate experiments were conducted, with each anthelmintic administered at increasing doses (6 in total) from 0.1 to 10,000 nM and compared to negative controls, such as 0.1% DMSO or water, as well as a positive control using 100 µg/mL doxycycline, an mitochondrial stress inducer that disrupts mitochondrial translation (47,48).

**Table 1:** Drugs and drug classes used in this study.

| Drug class | Drug name | Mode of action | References |
| --- | --- | --- | --- |
| **Benzimidazoles (BZ)** | Albendazole | β-tubulin inhibitor | (49) |
| **Macrocyclic lactones (MLs)** | Ivermectin<br>Milbemycin<br>oxime | Agonist of Glutamate and GABA-gated chloride channels | (50) |
| **Imidazothiazoles** | Levamisole | Agonist of acetylcholine receptors | (51) |
| **Amino-acetonitrile derivatives** | Monepantel | Agonist of acetylcholine receptors | (11) |
| **Cyclicoctadepsipeptides** | Emodepside | Agonist of SLO-1 receptors | (52) |
| **Diamidine derivative** | Tribendimidine | Modulator of acetylcholine receptors | (53) |
| **Salicylanilides** | Closantel | Disruption of energy metabolism – chitinase inhibitor | (54,55) |

The growth curves, generated by tracking the occupied area (µm²/h) of the worms during incubation, provided an overall perspective on the impact of each tested anthelmintic (Fig 1). From these initial results, several molecules and classes, including emodepside, macrocyclic lactones, and monepantel, stood out due to their growth inhibition at micromolar concentrations.

**Fig 1:**
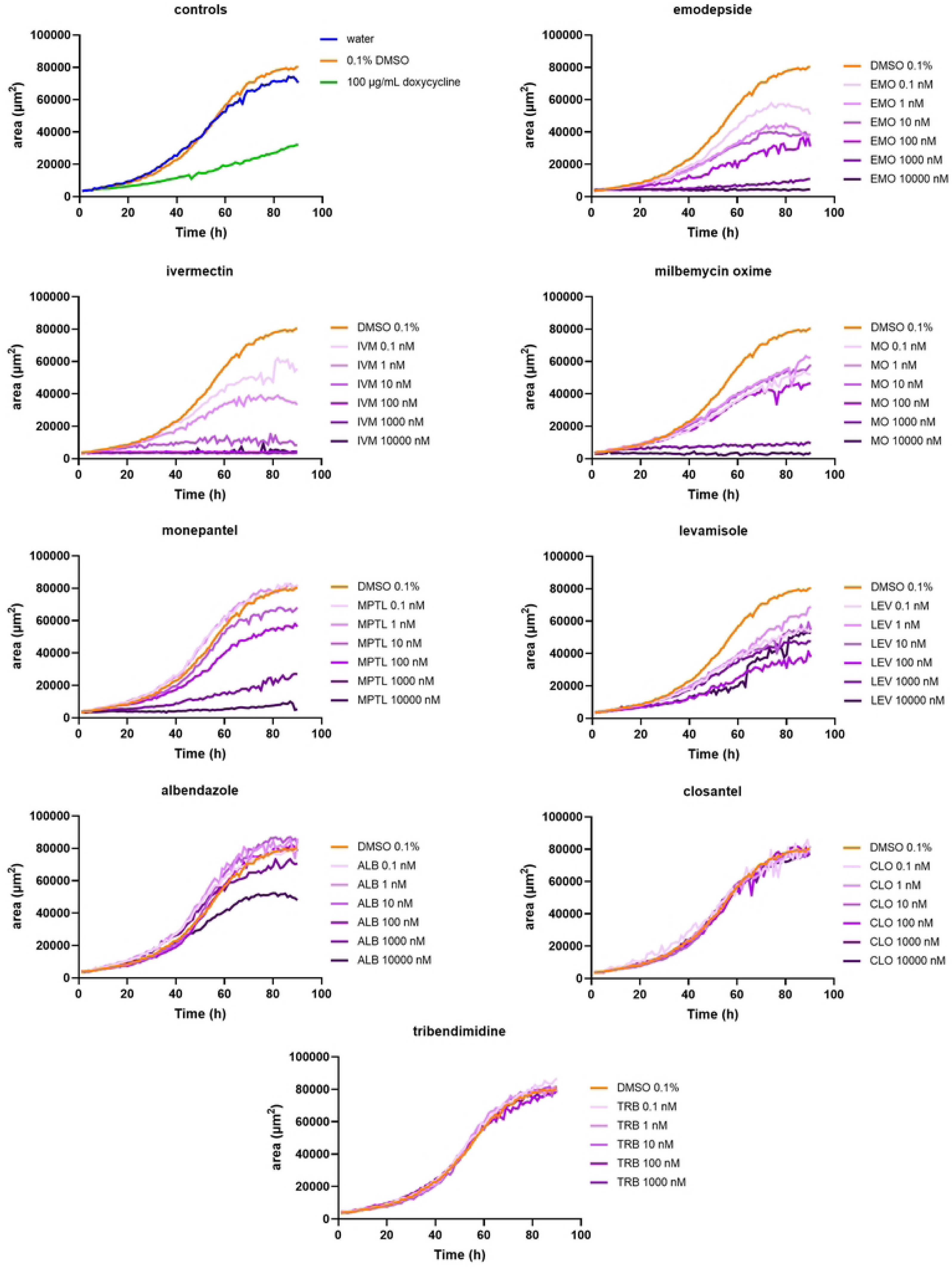
**Growth curves (area over time) of the SJ4100 strain for each treatment performed from the L1 stage to adult**. Average worm volume (area) was measured over time to assess the developmental progression of wild-type *C. elegans* exposed to various concentrations of anthelmintic compounds starting from the L1 larval stage. The orange curve represents vehicle-treated controls (0.1% DMSO), while the purple curves indicate responses to increasing drug concentrations. Data represent mean worm size at each time point, illustrating the impact of drug treatment on growth dynamics and developmental timing.

Conversely, the highest concentration (10 µM) of levamisole and albendazole resulted in areas under the curve of 65.3% and 59.7%, respectively, relative to the DMSO control (Fig 1), indicating a moderate effect on developmental speed. Finally, closantel and tribendimidine exhibited no significant effects on development, with growth curves and studied parameters closely aligning with DMSO control values (Fig 1).

Interestingly, parallel tests on N2 (wild-type) worms showed similar growth trends (S1 Fig). Slight differences were observed, with the transgenic strain being less impacted by levamisole than the N2 individuals, as only the top tested dose (10 µM) induced a significant reduction in the area under the curve (S1 Fig).

A series of images from each incubation chamber were taken throughout the test period from L1 larvae to adults. Representative images of each compound at the highest tested dose (10 µM) and after 90h of treatment are shown in Fig 2. At these levels, the image caption led to observe that the transgenic larval stages introduced into the system and exposed to emodepside, macrocyclic lactones, and monepantel were unable to develop, occupying minimal volume compared to the controls (Fig 2). The same visual inspection also led to notice albendazole treatment signatures in form of altered morphology and cuticular degradations while doxycycline prevented the individual from growing to the adult stage. Noteworthy, control experiments conducted on N2 strains did not reveal any major differences regarding the morphological features compared to the SJ4100 strain (S2 Fig). Additionally, no treatment condition, regardless of the dose, induced fluorescence emission indicative of mitochondrial stress response beyond the levels observed in negative control conditions. In contrast, doxycycline induced a GFP emission six times higher than that of the DMSO controls (data not shown).

**Fig 2:**
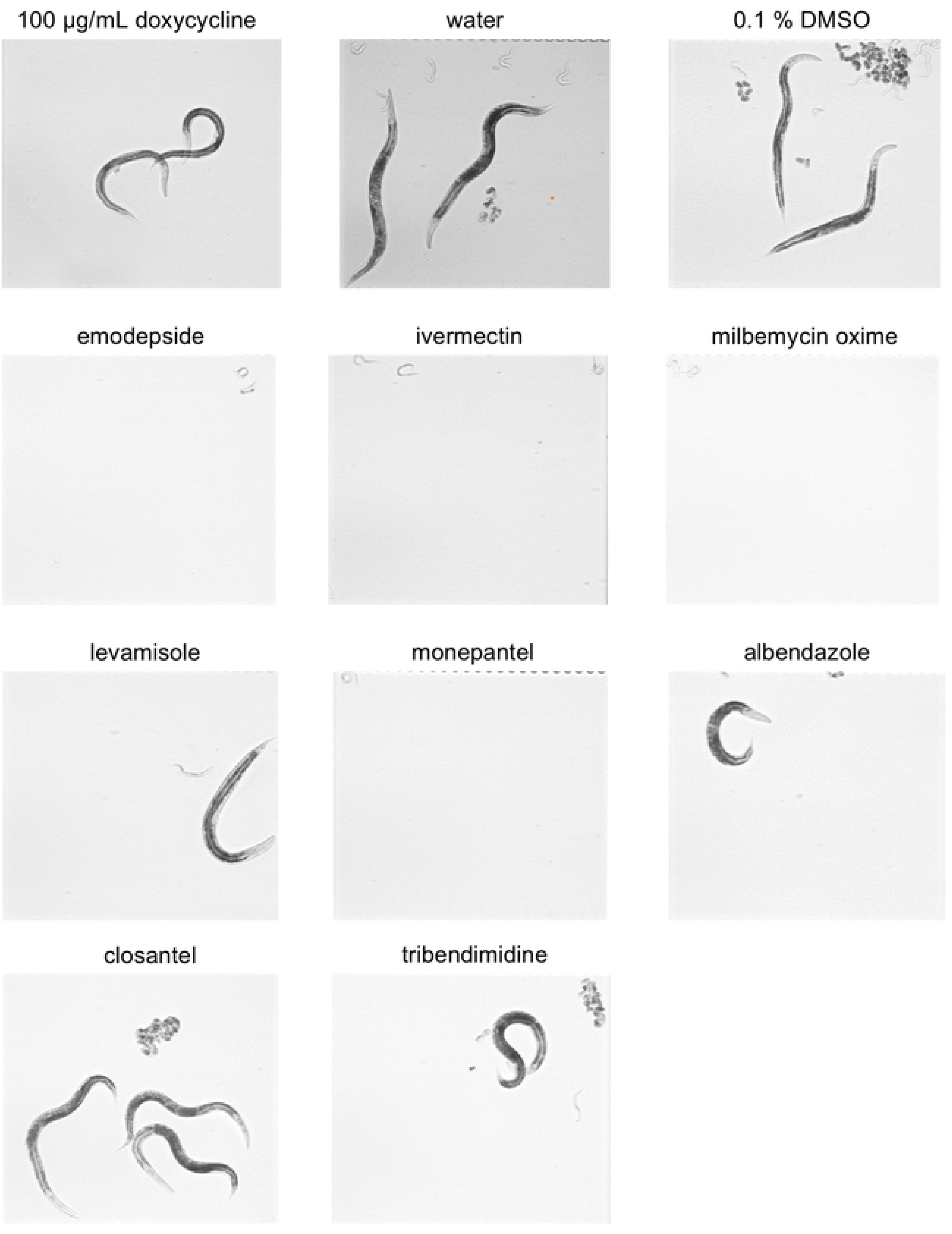
Light microscopic images showing SJ4100. *C. elegans* **treated with 10 µM of each anthelmintic.** Representative images of *C. elegans* after 10 µM anthelmintic exposure (90 hours). Each picture was taken through the SydLab™ integrated vision machinery using a 10X magnification and corresponds to one incubation chamber initially containing two to three L1 larvae. Controls consist of 0.1% DMSO, water or 100 µg/mL doxycycline treatment. The images highlight distinct morphological changes such as alterations in body shape, size, structure and progeny compared to the control group, emphasizing the differential impact of each anthelmintic on the nematode.

One of the numerous assets of SydLab™ system is also to run algorithms to quantify key aspects of worm development, such as worm volume (*K*), length (*K_length*), and surface area (*K_area*) at the experiment’s conclusion. Additionally, kinetic data, including the time needed for worms to reach half their final volume (*r*), length (*r_length*), and surface area (*r_area*), were recorded, enabling detection of any developmental delays or accelerations relative to controls (Table 2). In this context, treatments with increasing doses of emodepside, monepantel, or macrocyclic lactones significantly reduced the *K* parameters, with ratios dropping below 0.2 (emodepside) and 0.15 (ivermectin) at the highest concentration (10 µM) compared to the negative (DMSO) control (Fig 3). In contrast, doxycycline markedly slowed larval development, producing “r” ratios above 1 (Fig 3) while having minimal impact on worm dimensions (*K*). These findings suggest developmental arrest at the larval stage, preventing the expected morphological characteristics seen under negative control conditions. Analysis of the growth curves per treatment confirmed such a dose-dependent developmental slowdown for the compounds in question. Notably, even the lowest concentrations of emodepside and macrocyclic lactones (0.1 nM) did not produce growth curves similar to the negative control (Fig 1) and most importantly, developmental kinetics did not appear to be significantly affected. The identical development-based analysis performed in parallel onto the N2 strains confirmed the observation made with the SJ4100 strain (S3 Fig) with the most dramatic impact on worm development occurring with the aforementioned compounds.

**Fig 3:**
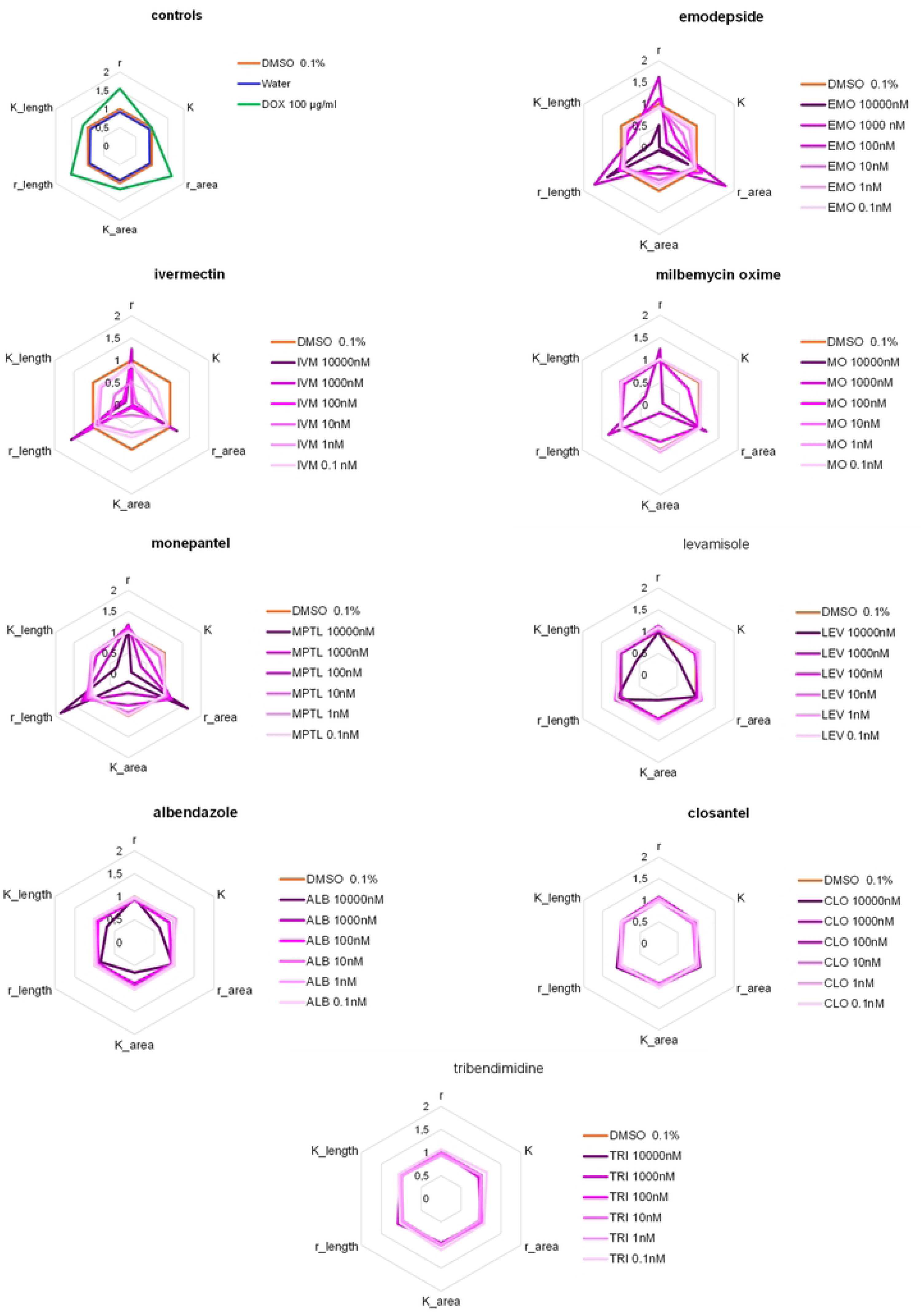
Radar plot illustrating the impact of anthelmintics on the development process of SJ4100 *C. elegans*, from the L1 larvae to the adult stage. Each axis represents a specific developmental feature monitored by the SydLab™ system. L1 larvae were exposed to six dilutions (purple lines) of each tested anthelmintic (10,000 to 0.1 nM) for a period of 95 hours until adult worms and subsequent progeny were obtained under the conditions of the negative control (0.1% DMSO, orange line). The plotted values in the graph indicate the average ratio calculated at the end of the assay from three replicated experiments compared to the negative control.

**Table 2:** Summary of parameters screened and registered by the SydLab™ system.

| Class | Parameter | Readout |
| --- | --- | --- |
| **development** | r | Time needed for the worms to reach half of their maximal volume |
|  | K | Maximal volumes of the worms |
|  | r area | Time needed for the worms to reach half of their maximal area |
|  | K area | Maximal are of the worms |
|  | r length | Time needed for the worms to reach half of their maximal length |
|  | K length | Maximal length of the worms |
| **morphology** | Shape 1 | Percentage of time that worms spend in straight shape (death/molting) |
|  | Shape 2 | Percentage of time that worms spend in active shape |
|  | Shape 3 | Percentage of time that worms spend in eating shape |
|  | Shape 4 | Percentage of time that worms spend in coiled shape (toxic environment) |
| **reproduction** | Fertility | 1 if eggs appearance (fertility); 0 if no eggs appearance (sterility) |
|  | Egg emergence | Time to which the first egg is detected |
|  | Embryotoxicity | 1 if eggs and progeny appearance (no embryotoxicity); 0 if eggs appearance but not progeny (embryotoxicity) |
|  | Progeny emergence | Time to which the first progeny is detected |
| **fluorescence** | AUC intensity | Area under the curve of the mean fluorescent intensity over time |

Taken as a whole, these data highlight the substantial influence of specific active compound classes, particularly emodepside, monepantel, and macrocyclic lactones, on developmental processes. In these cases, they induced larval arrest at early stages and severely affected worm size parameters.

### Anthelmintics induce recognizable morphological signatures that can be identified by machine vision

From a simple morphological perspective, anthelmintics create characteristic and dose-dependent signatures that are often related to their mode of action and targets (23,25,52,56). SydLab™ classifies these observations into four shapes: straight and non-motile (shape 1), active and motile (shape 2), feeding (shape 3), and coiled and/or curled (shape 4) (Table 2). Based on these morphological criteria, several anthelmintics, such as emodepside, levamisole, ivermectin, and especially monepantel, stand out by adopting a folded form (Figs 2 and 4 -5). This signature is typical, perfectly detectable by machine vision and particularly pronounced for monepantel and emodepside treatments, with a respective average ratio of 7.47 ± 3.12 and 3.77 ± 1.32 compared to negative controls. In contrast, albendazole resulted in fewer mobile worms with visibly degraded cuticles than the control worms (Fig 2). Finally, closantel and tribendimidine did not generate sufficient visible signatures for the algorithms, indicating a clear absence of morphological effects on exposed worms (Figs 2 and 4). Interestingly, comparing the transgenic strain to N2 worms has enabled recording noticeable discrepancies of emodepside, ivermectin and in some extents with levamisole treatments. Indeed, N2 worms exposed to these drugs were most recognizable by their shape 1 (straight and paralyzed) (S4 Fig and S5 A-B Figs), which contrasts, as described above, with transgenic worms primarily exhibiting shape 4 (coiled and/or curled). Both strains were severely impaired in terms of movement, with a noticeable larval arrest observed in both cases.

**Fig 4:**
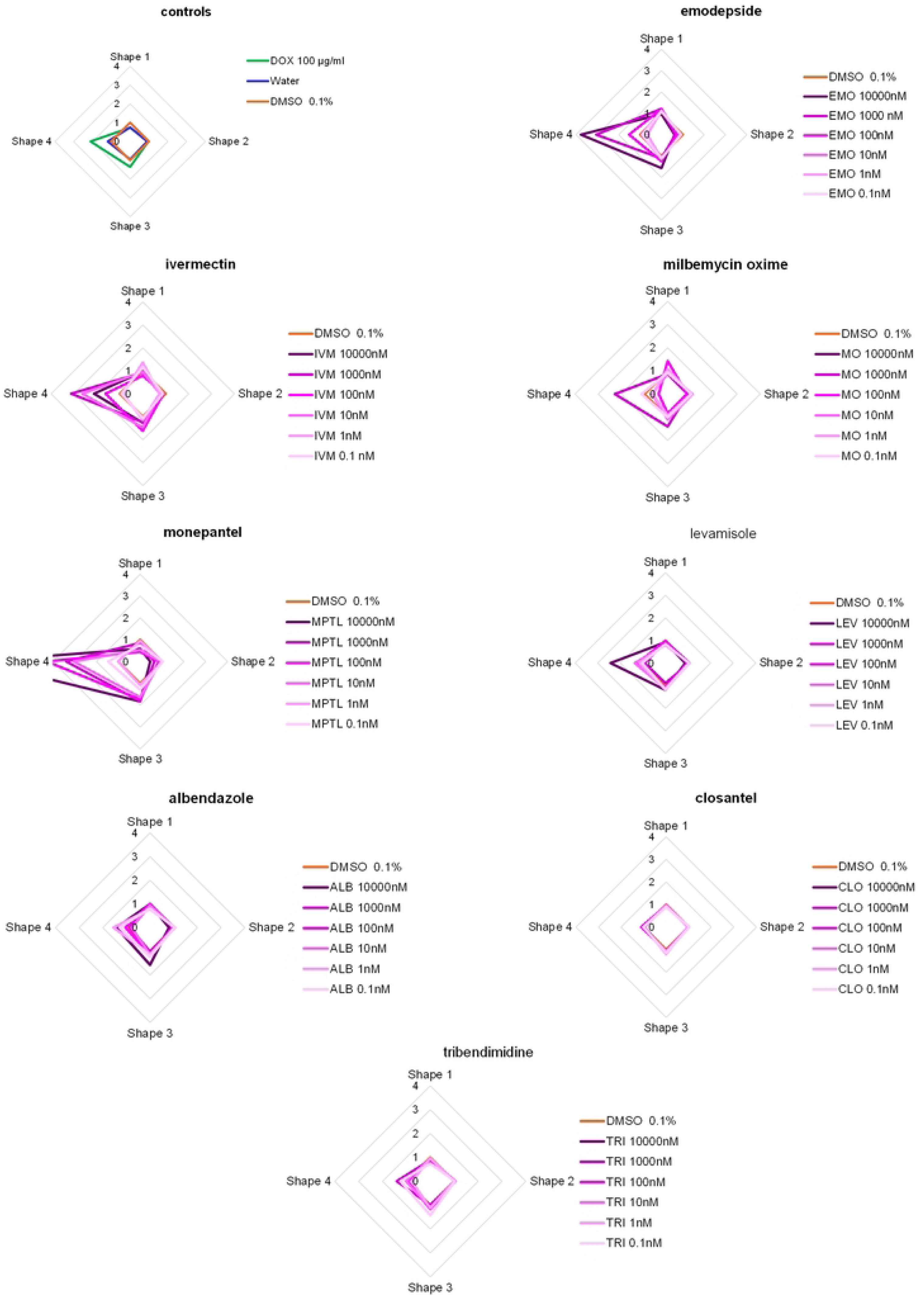
Radar plot illustrating the morphological impact of anthelmintics on SJ4100 *C. elegans*, from the L1 larvae to the adult stage. Each axis represents a specific shape feature (shape 1: stick-like, not moving; shape 2: active, in movement; shape 3: eating; shape 4: coiled) monitored by the SydLab™ system. L1 larvae were exposed to six dilutions (purple lines) of each tested anthelmintic (10,000 to 0.1 nM) for a period of 95 hours until adult worms and subsequent progeny were obtained under the conditions of the negative control (0.1% DMSO, orange line). The plotted values in the graph indicate the average ratio calculated at the end of the assay from three replicated experiments compared to the negative control.

**Fig 5.**
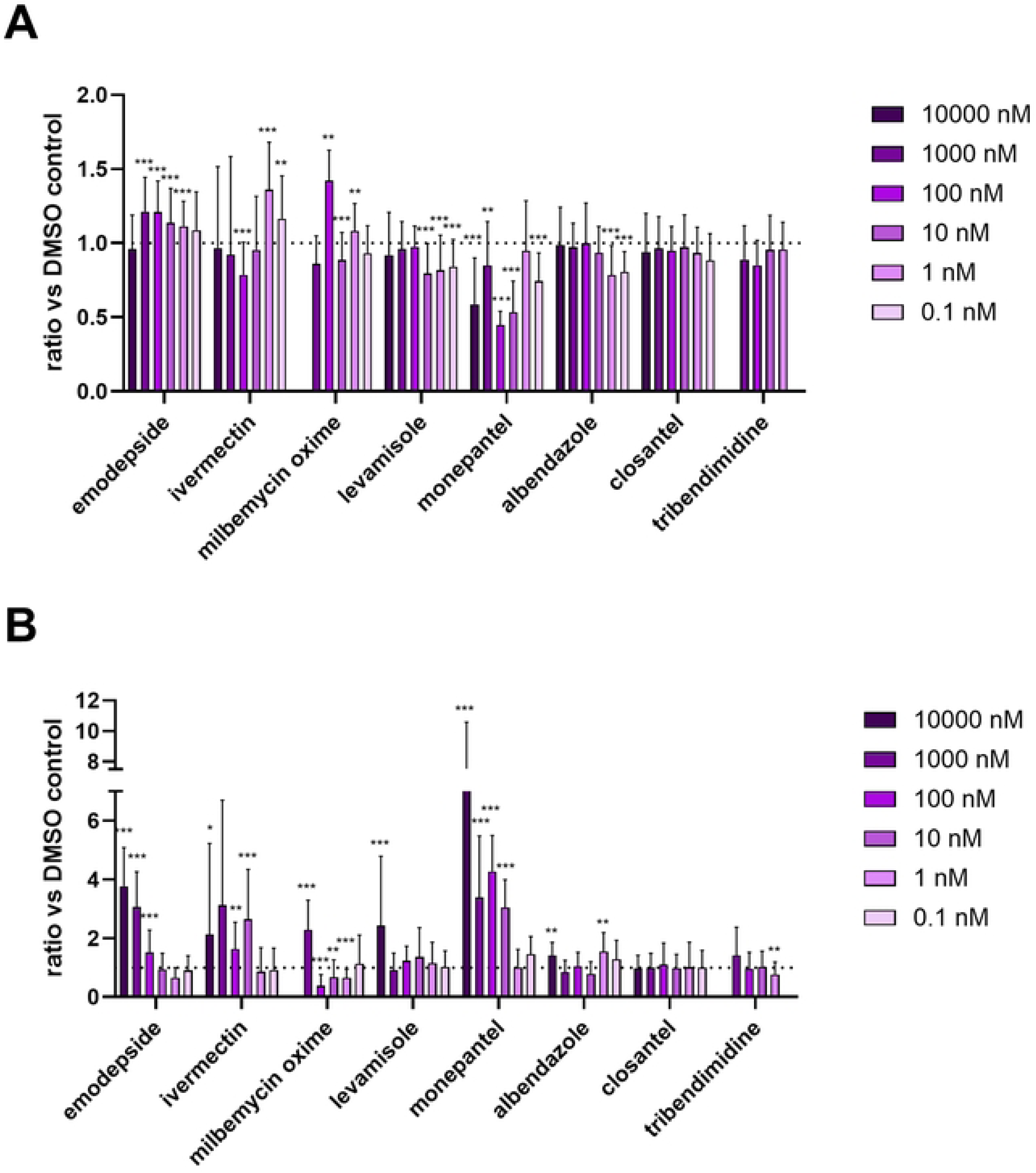
Dose-dependent quantification of shape 1 and 4 measured for each anthelmintic using the SJ4100 *C. elegans*, from the L1 larvae to the adult stage. Shape 1 (A) and 4 (B) ratios (calculated against DMSO values) were collected and calculated from at least three replicate experiments and plotted per anthelmintic ingredient and concentration. n per condition and t-test values are indicated in S1 Table. Dashed line is used as marker of ratio 1 reference. Asterisks indicate levels of statistical significance: *p ≤ 0.05, **p ≤ 0.005, ***p ≤ 0.0005. Comparisons with p-values greater than 0.05 are considered not statistically significant and are not marked with asterisks.

These results highlight the ability of SydLab™ to classify anthelmintic-induced morphological signatures, revealing distinct shape patterns correlated with drug mode of action. Additionally, strain-dependent differences in response to anthelmintics can be monitored and be processed in extended comparative analyses.

### Reproduction processes are diversely affected by anthelmintics upon constant exposure

One of the key capabilities of SydLab™ is to maintain worms under treatment with continuous food administration for an extended period. This allows for the prolonged development of the worms until they reach adulthood (if the treatment allows), and even until the emergence of eggs and larvae of the next generation. Regarding reproductive aspects (see Table 2 for details), a total of six additional parameters were studied, four of which are highlighted in Fig 6: fertility, time of first egg appearance, embryotoxicity of the treatments, and time of L1 larva emergence in the next generation. Worms exposed to 0.1% DMSO serve as the baseline for calculating ratios for each parameter.

**Fig 6:**
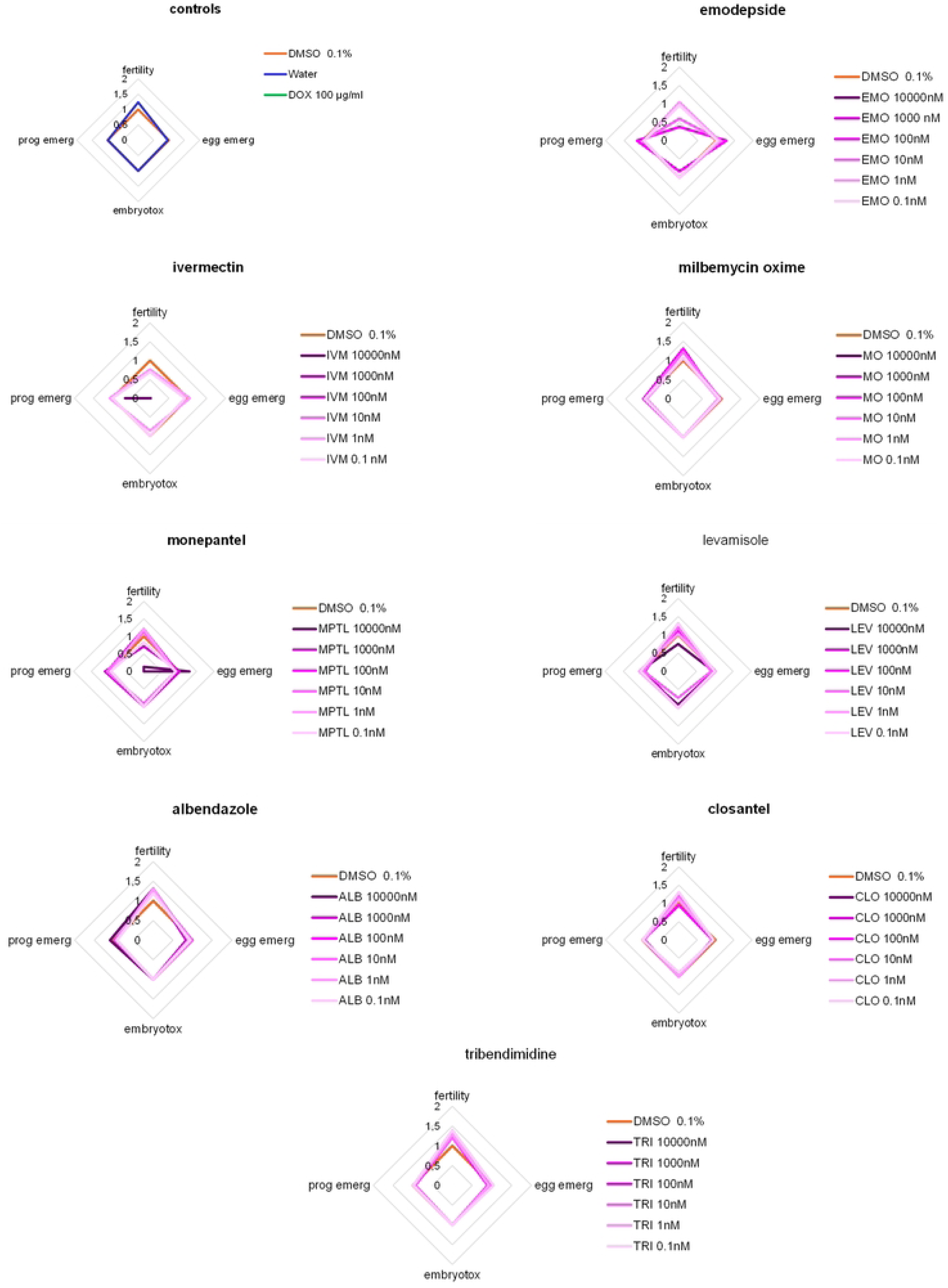
Radar plot illustrating the impact of anthelmintics on the main reproduction processes of SJ4100 *C. elegans*. Each axis represents a specific reproduction feature (see Table 2 for details) monitored by the SydLab™ system. L1 larvae were exposed to six dilutions (purple lines) of each tested anthelmintic (10,000 to 0.1 nM) for a period of 95 hours until adult worms and subsequent progeny were obtained under the conditions of the negative control (0.1% DMSO, orange line). The plotted values in the graph indicate the average ratio calculated at the end of the assay from three replicated experiments compared to the negative control.

Using this method, the larval developmental arrest observed earlier with certain anthelmintics carries over into reproductive analysis. For example, high doses of emodepside resulted in zero fertility (absence of eggs) starting at concentrations of 10 to 100 nM, which are the highest doses allowing the worms to reach adulthood (Fig 6). Concerning macrocyclic lactones, the algorithm is unable to assess the effects of doses above 10 nM (ivermectin) and 100 nM (milbemycin oxime) due to the persistent larval stages in the incubation chambers. However, for lower doses, no differences from control conditions were detected. Importantly, when comparing reproductive aspects between SJ4100 and N2 strains, no outlying observations could be made (S6 Fig). Regarding the other compounds, the highest dose of levamisole induces a reduction in fertility (ratio of 0.75 ± 0.66) but does not significantly affect the appearance of the few detected eggs or their progression to the L1 larval stage. Except for monepantel, none of the remaining analyzed molecules showed any embryotoxic or sterilizing effects.

These reproductive outcomes largely reflect either larval developmental arrest preventing worms from reaching maturity or the limited impact of certain drugs on the reproduction of adults that successfully developed.

### Validation of SydLab™ technology through comparative larval development and migration assays

Given that SydLab™ technology relies on complete automation of the *C. elegans* development process, it is important to compare the data obtained with those of a recognized classical test, whose standard protocol serves as a reference in numerous studies (3,19). Thus, SJ4100 strain was synchronized to initiate the developmental assay from the egg-to-L4 larval stage throughout a 48h incubation period. The motility of larvae exposed to each of the studied compounds at multiple doses was subsequently measured using a proprietary motility reading system (Invenesis Sàrl). The principle of this reading machine is based on motion detection imaging in a 384-well plate format.

Among the drugs tested in this protocol, several showed little to no effect on the motility of newly developed L4 larvae. Based on the analysis of the concentration-response curves (S7 Fig), this was particularly the case for levamisole, tribendimidine and closantel whose respective EC_50_ values exceeded 50 µM (Table 3) on the mutant strain. In contrast, macrocyclic lactones induced a complete lack of activity in the treated wells, with respective EC_50_ values of 0.01 ± 0.001 and 0.06 ± 0.003 µM for ivermectin and milbemycin oxime (Table 3). Similarly, neuromodulatory compounds such as emodepside and monepantel effectively inactivated the treated worms, with EC_50_ values of 0.23 ± 0.01 and 0.37 ± 0.03 µM, respectively. These additional data on larval development and final motility in the wells demonstrated a disparity in activity among the families of active compounds, consistent with previous observations using the SydLab™ system. Interestingly, the N2 strains was used in parallel, and several discrepancies of responses were identified. Indeed, EC_50_ calculation enabled identifying increased sensitivity to albendazole (5.5-fold) closantel (13.9-fold) with the wild-type worms while susceptibility to levamisole was again observable with a µm-ranged EC_50_ (11.83 ± 0.35 µM) observed at the tested dose-range (Table 3).

**Table 3:** Summary of effective concentrations collected from the *in vitro C. elegans* larval development assay.

| Compound | N2 Bristol | | SJ4100 (*hsp-6::gfp*) | | t-test<br>($p < 0.05$) |
| --- | --- | --- | --- | --- | --- |
|  | EC<sub>50</sub> (μM) | SE | EC<sub>50</sub> (μM) | SE |  |
| albendazole | 4,22 | 0,17 | 23,24 | 0,66 | 0,000010 (***) |
| ivermectin | 0,01 | 0,001 | 0,01 | 0,001 | 0,047421 (*) |
| milbemycin oxime | 0,15 | 0,001 | 0,06 | 0,003 | 0,000009 (***) |
| levamisole | 11,83 | 0,35 | > 100 | NA | NA |
| monepantel | 0,14 | 0,003 | 0,37 | 0,03 | 0,002370 (**) |
| emodepside | 0,25 | 0,03 | 0,23 | 0,01 | 0,698912 |
| tribendimidine | > 100 | NA | > 100 | NA | NA |
| closantel | 4,83 | 0,14 | 67,00 | NA | NA |

On a complementary aspect, motility is a fundamental phenotype for assessing the efficacy of most anthelmintics, as a large majority of them directly or indirectly target the functioning of the nematode neuromuscular system. However, during a classic immersion test lasting at least 48h, the endpoint motility measurement may not reflect the full potential of compounds, due to potential detoxification effects and other recovery phenomena. As an additional analysis, the migration trap assay, described in earlier studies (21,45,46), provides a valuable tool for assessing the motility and migratory behavior of parasitic larvae following a 24-hour exposure to anthelmintics. This method is particularly useful for evaluating the neuromuscular effects of drugs like ivermectin, which inhibit locomotion by targeting glutamate-gated chloride channels in the worms, effectively trapping them in the migration assay (45). The use of this complementary technique ensures a more comprehensive assessment of the drug’s efficacy, beyond simple motility measurements.

L3 larvae from the transgenic strain were freshly cultured and exposed to increasing concentrations of each active molecule. After this treatment step, the migration of the worms was observed and recorded after a period defined by the worms treated with the DMSO control. In this way, dose-response curves were generated (S8 Fig), and EC_50_ values were calculated (Table 4). Among the tested anthelmintics, tribendimidine and closantel did not produce any deleterious effects on the migration process, making the calculation of EC_50_ inapplicable. The least effective molecules were albendazole (EC_50_ = 27.21 ± 6.87 µM) and levamisole (EC_50_ = 12.13 ± 0.47 µM), followed by emodepside (EC_50_ = 8.79 ± 0.36 µM). In contrast, macrocyclic lactones were detected at relatively low doses with calculated EC_50_ values of 0.18 ± 0.01 and 0.68 ± 0.03 µM for ivermectin and milbemycin oxime, respectively. It is noteworthy that monepantel also acts very effectively on migrating adults, with its effectiveness levels detected in the nanomolar range (EC_50_ = 0.50 ± 0.01 µM).

**Table 4:** Summary of effective concentrations collected from the *in vitro C. elegans* migration trap assay.

| Compound | N2 Bristol | | SJ4100 (*hsp-6::gfp*) | | t-test ($p < 0.05$) |
| --- | --- | --- | --- | --- | --- |
|  | EC<sub>50</sub> (μM) | SE | EC<sub>50</sub> (μM) | SE |  |
| **albendazole** | **20,44** | 1,36 | **27,21** | 6,87 | 0,389078 |
| **ivermectin** | **0,04** | 0,00 | **0,18** | 0,01 | 0,000151 (***) |
| **milbemycin oxime** | **0,40** | 0,01 | **0,68** | 0,03 | 0,000898 (**) |
| **levamisole** | **13,14** | 0,42 | **12,13** | 0,47 | 0,187813 |
| **monepantel** | **0,28** | 0,01 | **0,50** | 0,01 | 0,000100 (***) |
| **emodepside** | **4,40** | 0,09 | **8,79** | 0,36 | 0,000295 |
| **tribendimidine** | **NA** | NA | **1,85** | 0,31 | NA |
| **closantel** | **2,49** | 0,05 | **NA** | NA | NA |

The MTA revealed significant differences in compound sensitivity between the N2 and SJ4100 strains for several anthelmintics. Notably, SJ4100 worms exhibited reduced sensitivity to ivermectin, milbemycin oxime, monepantel, and emodepside, as indicated by significantly higher EC_50_ values compared to N2 (p < 0.001 for ivermectin and monepantel; p < 0.005 for milbemycin; p < 0.001 for emodepside). In contrast, no significant differences in EC_50_ were observed between strains for albendazole and levamisole, suggesting similar responses to these compounds. Comparative analysis remained limited with tribendimidine and closantel between strains due to an ineffective dose-range leading to hardly computable EC_50_ values (Table 4).

Altogether, these data highlight the level of sensitivity achieved when motility was challenged. These data provide a broader perspective on the capacities of the compound and offer insights into their efficacy at doses higher than those tested by SydLab™, while also exploring complementary biological processes induced by the treatments and throughout different incubation periods.

Freshly extracted eggs from both strains were exposed to 10 serial dilutions of each compound, starting at 100 µM with 3.16-fold dilution steps. After 48h incubation, motility was recorded and normalized efficacies were computed (S3 Table). Then, concentration-efficacy curves were generated from triplicate measurements and EC_50_ values (µM) were calculated by fitting the data to a four-parameter logistic (4PL) model constrained between 0% and 100% efficacy. The mean EC_50_, and standard error of the mean (SE) are reported for each compound. Statistical comparison of EC_50_ values between the N2 and SJ4100 strains was performed using multiple unpaired t-tests (GraphPad Prism, version 10), with p < 0.05 considered statistically significant. (*) p ≤ 0.05, (**) p ≤ 0.005, (***) p ≤ 0.0005. Comparisons with p-values greater than 0.05 are considered not statistically significant and are not marked with asterisks.

Freshly cultivated L3s from both strains were exposed to 10 serial dilutions of each compound, starting at 100 µM with 3.16-fold dilution steps. After 24h incubation, worms were deposited in the MTA device, motility was recorded and normalized efficacies were computed (S3 Table). Then, concentration-efficacy curves were generated from triplicate measurements and EC_50_ values (µM) were calculated by fitting the data to a four-parameter logistic (4PL) model constrained between 0% and 100% efficacy. The mean EC_50_, and standard error of the mean (SE) are reported for each compound. Statistical comparison of EC_50_ values between the N2 and SJ4100 strains was performed using multiple unpaired t-tests (GraphPad Prism, version 10), with p < 0.05 considered statistically significant. (*) p ≤ 0.05, (**) p ≤ 0.005, (***) p ≤ 0.0005. Comparisons with p-values greater than 0.05 are considered not statistically significant and are not marked with asterisks.

## Discussion

This study explored the effects of eight anthelmintics on *Caenorhabditis elegans* development, motility, reproduction, and morphology using the SydLab™ platform, with additional insights collected from a non-robotic larval development and migration assays. While our findings confirm dose-dependent effects of some of these compounds, the choice of a dose range with an upper limit of 10 µM was determined as a compromise between the need to observe the effects of molecules with varying levels of efficacy and solubility (e.g., the inability to test tribendimidine at 10 µM) and cost-effectiveness in the context of screening campaigns. Moreover, this concentration was chosen to standardize the screening conditions across all eight anthelmintics, as this is a commonly applied concentration in high-throughput screening (HTS) strategies (31) as compounds exhibiting activities below this threshold serve as robust leads for subsequent drug development. In this study, the limitations of the top dose used may have influenced the ability to fully capture the efficacy of certain anthelmintics, to capture more obvious phenotypic signatures that would lead to a better understanding of particular modes of action. For example, albendazole, a widely used broad-spectrum anthelmintic, has been shown to require higher dose ranges to achieve significant effects in nematode models due to its relatively low water solubility and the need for prolonged exposure to reach sufficient intracellular concentrations (49). In studies on parasitic nematodes such as *Haemonchus contortus*, albendazole exhibited significant anthelmintic activity only at doses ranging from 20 µM to 50 µM. Similarly, closantel, another compound tested in this study, is known for its activity against trematodes and cestodes, but its efficacy rate on nematodes is notably challenged by differences in its mode of action and its reduced ability to penetrate nematode cuticle barriers (57). In our study, closantel did not produce significant developmental or motility effects at 10 µM, suggesting that higher doses would be needed to observe its full potential against nematodes.

In terms of developmental effects, compounds like emodepside, monepantel, and macrocyclic lactones showed a marked impact on larval growth. The SydLab™ system tracked reductions in worm size, surface area, and volume, particularly at the highest doses. These effects are consistent with previous studies showing that emodepside and macrocyclic lactones arrest larval development through disruption of neuromuscular communication (25,58). For instance, emodepside caused a significant reduction in larval size at doses as low as 1 µM, in agreement with other reports where emodepside induced arrest at even lower concentrations (e.g., 0.5 µM) in other nematode models (25). Likewise, ivermectin and milbemycin oxime induced strong growth inhibition, with larvae failing to progress beyond early larval stages at concentrations starting from 0.1 µM, which is consistent with their mechanism of action on chloride channels (59). In contrast, levamisole and albendazole showed only moderate developmental effects at 10 µM, with worms retaining partial growth compared to controls, a finding that aligns with previous research showing that these compounds can require higher concentrations to exert stronger developmental arrest (23,49). When comparing developmental effects between the SJ4100 and N2 strains, no major discrepancies were observed, likely because both strains share conserved developmental pathways targeted by these compounds. However, slight differences were noted for levamisole, as N2 worms were less impacted than transgenic individuals. This could be attributed to the genetic background of the genetically modified worms, which express oxidative stress-related fluorescence markers that may indirectly influence sensitivity to certain compounds, although no significant signs of stress could be detected with any of the tested treatments. Nevertheless, these differences were only significant at the highest dose tested (10 µM; Fig 1 and S1 Fig).

The effects on reproduction also offered variable spectrum of responses. SydLab™ allowed for the continuous tracking of reproductive processes, such as fertility and time of egg laying. At high doses, emodepside completely suppressed egg production, leading to zero fertility, an effect reported in other studies where emodepside blocked reproduction by disrupting motor neurons that control egg laying (52). Monepantel also caused a significant reduction in fertility, with fewer eggs detected at concentrations as low as 1 µM. The absence of eggs in these cases suggests that the compounds induce a form of reproductive arrest that prevents the transition to adulthood, similar to findings in parasitic nematodes like *Teladorsagia circumcincta* (11). Conversely, compounds like albendazole, levamisole, and tribendimidine had less pronounced effects on fertility, which may be linked to their weaker impact on development at the doses tested. Importantly, no significant differences in reproductive parameters were observed between the transgenic and N2 strains (Fig 6 and S6 Fig). This is likely due to the fact that the reproductive processes in *C. elegans* rely on highly conserved neuromuscular and hormonal pathways that are equally targeted by these compounds in both strains. Additionally, the effects on reproduction appear to depend more on the developmental disruption induced by the compounds rather than any intrinsic strain-specific sensitivity.

The morphological effects observed, particularly in worms treated with emodepside, levamisole, and ivermectin, were distinctive. SydLab™ was able to classify worms into different shapes, including coiled and folded forms, which are indicative of neuromuscular disruption. Emodepside and monepantel produced characteristic folded worms, consistent with paralysis or impaired locomotion resulting from interference with neurotransmission. Levamisole, a nicotinic acetylcholine receptor agonist, induced coiling in a dose-dependent manner, a phenotype that has been linked to the overactivation of muscle contractions (60). Interestingly, comparing transgenic and N2 strains revealed notable morphological differences for certain compounds, particularly emodepside and ivermectin. N2 worms were predominantly observed in a straight, paralyzed form (shape 1), while SJ4100 worms exhibited more coiled and curled forms (shape 4) (Fig 5, S5 Fig). These differences may stem from the genetic background of transgenic worms, which can exacerbate variability of neuromuscular disruptions caused by these compounds. Indeed, natural genetic variation plays a crucial role in shaping nematode responses to emodepside. Wit et al. (61) demonstrated that wild *C. elegans* strains exhibit diverse sensitivities to emodepside, with observations identifying a hormetic effect where low emodepside concentrations increased brood size. Such findings highlight the importance of considering both genetic background and dosing strategies when evaluating emodepside efficacy and subsequent phenotypic effects. Despite these morphological differences, both strains were similarly impaired in terms of motility and exhibited larval arrest under these treatments. This indicates that while the genetic modification of SJ4100 worms may amplify certain phenotypic responses, the underlying effects of these compounds on essential neuromuscular and developmental pathways are conserved across strains. The ability of SydLab™ to capture these distinct morphological changes through high-resolution imaging underscores the platform’s strength in phenotypic screening.

The cuticle barrier of *C. elegans* likely played a role in limiting the efficacy of certain compounds, particularly those that are less lipophilic, such as albendazole and closantel. As noted in previous studies (24), the cuticle serves as a selective barrier that can impede the uptake of larger or less permeable compounds, reducing their intracellular efficacy. The composition of the cuticle, rich in collagen proteins and glycoproteins, creates a physical and chemical shield that many anthelmintics struggle to penetrate (32). This may explain why the activity of these compounds appeared limited at 10 µM, while more lipophilic compounds, such as emodepside and macrocyclic lactones, were able to bypass this barrier and exert their effects at much lower doses.

The larval development assay (LDA) and migration trap assay (MTA) provided important complementary information to the SydLab™ data. The LDA was particularly useful for confirming developmental arrest in larvae exposed to macrocyclic lactones, monepantel, and emodepside, reinforcing the notion that these compounds target key developmental pathways through the binding of receptors already targetable in the larval stages (19). However, the LDA’s focus on continuous exposure and growth endpoints may miss the transient effects seen with neuromodulatory compounds like levamisole, where recovery of motility after initial paralysis can mask developmental delays. In contrast, the MTA offered more granular insights into the motility effects of anthelmintics. By assessing the ability of larvae to migrate after drug exposure, the MTA effectively captured the neuromuscular impact of drugs like ivermectin and monepantel, which showed potent inhibition of migration at sub micromolar concentrations (45). The MTA’s focus on motility, independent of developmental endpoints, allows for a more direct evaluation of how these compounds disrupt neuromuscular function, making it a valuable tool alongside the SydLab™ system (21).

One of the key strengths of the SydLab™ platform is its ability to simultaneously track multiple phenotypic parameters, such as growth, morphology, and motility, across different concentrations and over extended periods. Additionally, SydLab™ has the capability to analyze multiple *C. elegans* strains at once, including genetically modified strains that reveal insights into specific biological pathways. In this study, the use of the SJ4100 strain allowed us to monitor mitochondrial stress response levels in response to anthelmintic treatments. Looking forward, the application of SydLab™ directly to parasitic species could greatly expand its utility, offering a more direct evaluation of anthelmintic efficacy and resistance mechanisms in species of veterinary and clinical relevance. This would also allow for a more detailed exploration of the modes of action of different anthelmintics and how they interact with specific parasitic systems.

While the 10 µM top dose used in this study may not have been sufficient to fully capture the efficacy of certain anthelmintics, the combination of SydLab™, LDA, and MTA provided a robust and complementary framework for evaluating drug effects on *C. elegans*. The results highlight the importance of using multiple assays to capture both developmental and motility-related phenotypes, particularly for compounds with diverse modes of action. Future studies should consider testing higher dose ranges for compounds like albendazole and closantel and expanding the application of SydLab™ to parasitic species to further elucidate anthelmintic mechanisms.

## Materials and methods

### *C. elegans* strains and maintenance

N2 wild-type and *hsp-6::gfp* (SJ4100) transgenic strains were provided by the *Caenorhabditis* Genetics Center, University of Minnesota, Minneapolis, MN, USA. The stock of worms for both strains was maintained by serially passaging the worms onto solid nematode growth medium (NGM) agar plates that were previously seeded with *Escherichia coli* OP50.

### SydLab™ one recordings

The effect of anthelmintics on larval development, reproduction, morphology, growth kinetics, and oxidative stress in *C. elegans* was studied using the SydLab™ system (Nagi Bioscience SA, Switzerland). This machine was fully equipped with a microfluidic-based platform and automated data recording and analysis software as previously described (Cornaglia et al., 2015; Preston et al., 2017; Atakan et al., 2019).

Prior to starting SydLab™, worms were synchronized to the L1 larval stage in complete S-medium. Briefly, young adult worms were collected by filtration and incubated overnight in complete S-medium. The freshly arrested L1 larvae were separated from the adults by filtration, obtaining a synchronized L1 larvae population and diluted in complete S-medium (S-basal solution (100 mM NaCl; 58 mM K_2_HPO_4_; 44 mM KH_2_PO_4_ 5mg/mL of cholesterol) supplemented with 3 mM CaCl_2_; 3 mM MgSO_4_; 10 mM potassium citrate; trace metal solution of EDTA, FeSO_4_, MnCl_2_, ZnSO_4_, CuSO_4_ ; 50 µg/mL carbenicillin; 50 µg/mL ampicillin; 0.5 µg/mL doxycycline and 0.06% Tween 20. L1 larvae were then injected into the microfluidic chip and exposed for a total period of 120h to six concentrations (0.01 to 10,000 nM, 10-fold dilutions) of each of the following anthelmintics: ivermectin, milbemycin oxime, levamisole, emodepside, closantel, albendazole (Sigma-Aldrich, USA), monepantel (MedChemExpress, USA) except for tribendimidine (MedChemExpress, USA) for which worms were exposed to five concentrations only (0.01 to 1,000 nM, 10-fold dilutions). Worms were fed with freeze-dried *E. coli* OP50 prepared in complete S-medium as described above. SydLab™ captured bright-field images of each incubation chamber every hour for the entire 120h assay duration, allowing time-dependent growth analysis.

Growth kinetics were calculated using the SydLab™ software. Briefly, image sets were analyzed using software algorithms developed by Nagi Bioscience SA, allowing to extract data about the size of each worm. From the model used to describe the growth, the software algorithms extracted two parameters: K, which is the maximum size (volume, area and length) the worm is aiming at, and r, which corresponds to the time when the worm reaches its half-size and determines the dynamic of the growth. This model is fitted to data with the Markov chain Monte Carlo (MCMC) method. For egg laying worms, the progeny was computationally removed (selected through their size) so fits are not altered. For all the fits, a threshold was applied to the Pearson’s chi-squared test, which measures the goodness of fit, to only keep the sensible ones.

To measure the mitochondrial stress response, SJ4100 worms were loaded into the microfluidic platform following the same principle. Bright-field and fluorescent images were also captured every hour for the entire assay duration, and the fluorescence intensity (GFP) was quantified using the SydLab^TM^ software to measure the induction of the mitochondrial stress response.

### *In vitro* larval development assay

N2 and SJ4100 worm strains were allowed to grow separately until the adult stage. Gravid adults were collected through several washing steps of NGM plates with cold M9 buffer (22 mM KH_2_PO_4_, 42 mM Na_2_HPO_4_, 86 mM NaCl, and 1 mM MgSO_4_) and transferred to a 50 mL tube. Adult worms were pelleted by centrifuging at 2000 g for five minutes at 10°C. The supernatant was then pipetted out to leave only 3.5 mL of the medium. 1.5 mL of a solution of NaOCl and 0.5 mL 5M NaOH were added to the worm solution. Worm lysis was carefully monitored over a maximum period of 8 min with continuous stirring. When no trace of dead adult bodies was visible, cold sterile distilled water was poured up to a volume of 50 mL and tubes were immediately centrifuged at 1500 g for 1 min (10°C). The maximum volume of liquid was removed manually, and two additional washing steps were performed with water, followed by a final liquid change with M9 buffer. The last centrifugation step removed most of the supernatant, leaving 5 mL of the egg solution. Eggs were counted and the volume was adjusted to reach a final concentration of 33 eggs/µL.

In the meantime, for the day of the experiment, 10 mL of OP50 liquid culture (Luria Broth medium with no antibiotics) was grown overnight at 37°C under gentle shaking. Bacteria were then diluted to 66% v/v in 0.18% NaCl to serve as a developmental medium for worms. For a single 384-well plate, a total volume of 8 mL (eggs with bacteria) was required, which implied mixing 1.85 mL of egg solution previously prepared and 6.15 mL of diluted bacteria.

A white clear-bottom (low volume) 384-well plate (Greiner, Germany) was filled with 5 µL/well of anthelmintic dilutions (100 to 10^-7^ µM in 10-fold steps), or controls, and 11 µL of egg/bacterial solution was automatically dispensed using a multidrop 384 ThermoLabsystem (ThermoScientific, USA). The plates were then sealed using Easyseal™ multiwell plate sealers (Merck KGaA, Germany). For motility measurement, newly emerged L4 larvae were recorded after 48h (N2) or 96h (SJ4100) using the proprietary Invenesis Sàrl machine vision system, which tracked pixel displacement to calculate motility as a percentage of the DMSO-treated controls.

### Migration trap assay

The Invenesis migration trap assay (MTA), previously described (45,46), was used to measure the effect of anthelmintics on the larval stages of *C. elegans* (N2 and SJ4100). Approximately 250 L3s were exposed to the drug or control treatments in each well of a 96-well plate. In all experiments, the larvae were exposed for 24 h to the compound in a solution of 1.5% DMSO and 0.00425% Tween-20. The larvae were then transferred to a migration plate, which is a 96-well plate allowing them to migrate from a deposit area to a trap area through an artificial corridor. Worm migration was monitored within a defined time window—based on DMSO controls—using an automated imaging system (equipped with a Basler acA2000-50 gm camera) measuring pixel displacement. The effect of compounds was expressed as a percentage reduction in motility compared to negative controls. Anthelmintics were tested in triplicate in 10 doses starting from 100 µM with 3.16-fold dilutions.

### Data analysis and statistics

Statistical analysis was performed using GraphPad Prism version 10 version (GraphPad Software, San Diego, California, USA). EC₅₀ values were calculated using nonlinear regression models (four-parameter logistic curves), fitted to dose-response data.

For the SydLab™ based screening, derived parameters (r, K, shapes and reproduction) were normalized to vehicle controls (0.1% DMSO), and the ratio of treatment to control was computed for each condition. Differences between each drug condition and the control were analyzed using unpaired two-tailed t-tests. A threshold approach was used to interpret biological significance: treatments were classified as “TOXIC,” “NEGATIVE,” or “ACUTE” based on the response ratio relative to predefined bounds. Statistical significance was defined at *p* < 0.05.

For the LDA and MTA datasets, raw readouts (motility or migration) were converted to efficacy scores (percent reduction compared to controls), and median efficacy was calculated per condition. No imputation was performed for missing values. Where replicates were available, comparisons between groups (e.g., WT *vs.* SJ4100 or different concentrations) were evaluated using unpaired two-tailed t-tests. Normality was assumed, and variance homogeneity was considered. Results are reported with median and raw data to highlight intra-group variability.

## Supporting information

**S1 Fig: Growth curves (area over time) of the N2 strain for each treatment performed from the L1 stage to adult.** Average worm volume (area) was measured over time to assess the developmental progression of wild-type *C. elegans* exposed to various concentrations of anthelmintic compounds starting from the L1 larval stage. The orange curve represents vehicle-treated controls (0.1% DMSO), while the purple curves indicate responses to increasing drug concentrations. Data represent mean worm size at each time point, illustrating the impact of drug treatment on growth dynamics and developmental timing.

**S2 Fig: Light microscopic images showing N2 *C. elegans* treated with 10 µM of each anthelmintic.** Representative images of *C. elegans* after 10 µM anthelmintic exposure (90 hours). Each picture was taken through the SydLab™ integrated vision machinery using a 10X magnification and corresponds to one incubation chamber initially containing two to three L1 larvae. Controls consist of 0.1% DMSO, water or 100 µg/mL doxycycline treatment. The images highlight distinct morphological changes such as alterations in body shape, size, structure and progeny compared to the control group, emphasizing the differential impact of each anthelmintic on the nematode.

**S3 Fig: Radar plot illustrating the impact of anthelmintics on the development process of N2 *C. elegans*, from the L1 larvae to the adult stage.** Each axis represents a specific developmental feature monitored by the SydLab™ system. L1 larvae were exposed to six dilutions (purple lines) of each tested anthelmintic (10,000 to 0.1 nM) for a period of 95 hours until adult worms and subsequent progeny were obtained under the conditions of the negative control (0.1% DMSO, orange line). The plotted values in the graph indicate the average ratio calculated at the end of the assay from three replicated experiments compared to the negative control.

**S4 Fig: Radar plot illustrating the morphological impact of anthelmintics on N2 *C. elegans*, from the L1 larvae to the adult stage.** Each axis represents a specific shape feature (shape 1: stick-like, not moving; shape 2: active, in movement; shape 3: eating; shape 4: coiled) monitored by the SydLab™ system. L1 larvae were exposed to six dilutions (purple lines) of each tested anthelmintic (10,000 to 0.1 nM) for a period of 95 hours until adult worms and subsequent progeny were obtained under the conditions of the negative control (0.1% DMSO, orange line). The plotted values in the graph indicate the average ratio calculated at the end of the assay from three replicated experiments compared to the negative control.

**S5 Fig: Dose-dependent quantification of shape 1 and 4 measured for each anthelmintic using the N2 *C. elegans*, from the L1 larvae to the adult stage.** Shape 1 (A) and 4 (B) ratios (calculated against DMSO values) were collected and calculated from at least three replicate experiments and plotted per anthelmintic ingredient and concentration. n per condition and t-test values are indicated in S2 Table. Dashed line is used as marker of ratio 1 reference. Asterisks indicate levels of statistical significance: *p ≤ 0.05, **p ≤ 0.005, ***p ≤ 0.0005. Comparisons with p-values greater than 0.05 are considered not statistically significant and are not marked with asterisks.

**S6 Fig: Radar plot illustrating the impact of anthelmintics on the main reproduction processes of N2 *C. elegans*.** Each axis represents a specific reproduction feature (see Table 2 for details) monitored by the SydLab™ system. L1 larvae were exposed to six dilutions (purple lines) of each tested anthelmintic (10,000 to 0.1 nM) for a period of 95 hours until adult worms and subsequent progeny were obtained under the conditions of the negative control (0.1% DMSO, orange line). The plotted values in the graph indicate the average ratio calculated at the end of the assay from three replicated experiments compared to the negative control.

**S7 Fig: Concentration-efficacy curves collected from the *in vitro C. elegans* larval development assay.** Freshly extracted eggs from both strains were exposed to 10 serial dilutions of each compound, starting at 100 µM with 3.16-fold dilution steps. After 48h incubation, motility was recorded and normalized efficacies were computed (S3 Table). Then, concentration-efficacy curves were generated from triplicate measurements and EC_50_ values (µM) were calculated by fitting the data to a four-parameter logistic (4PL) model constrained between 0% and 100% efficacy.

**S8 Fig: Concentration-efficacy curves collected from the *in vitro C. elegans* migration trap assay.** Freshly cultivated L3s from both strains were exposed to 10 serial dilutions of each compound, starting at 100 µM with 3.16-fold dilution steps. After 24h incubation, worms were deposited in the MTA device, motility was recorded and normalized efficacies were computed (S3 Table). Then, concentration-efficacy curves were generated from triplicate measurements and EC_50_ values (µM) were calculated by fitting the data to a four-parameter logistic (4PL) model constrained between 0% and 100% efficacy. Red values indicate the mean EC_50_.

**S1 Table: Summary of data collected from the SydLab™ using the SJ4100 *C. elegans* strain.** This table presents raw and normalized data related to developmental, morphological, and reproductive phenotypes measured in *C. elegans* expressing the mitochondrial stress reporter *hsp-6::gfp*. Animals were exposed to various anthelmintic compounds (10 μM) or vehicle control. Each sheet corresponds to a specific compound and includes parameters such as raw fluorescence intensity, calculated metrics (e.g., r and K), ratio to control, and number of animals analyzed. The dataset also includes results of statistical analyses (t-tests), thresholds for toxicity classification, and annotations regarding compound toxicity (e.g., TOXIC, NEGATIVE, ACUTE). This comprehensive dataset supports the characterization of drug-induced developmental and physiological perturbations *in vivo*.

**S2 Table: Summary of data collected from the SydLab™ using the N2 (wild-type) *C. elegans* strain.** This table presents raw and normalized data related to developmental, morphological, and reproductive phenotypes measured in wild-type *C. elegans*. Animals were exposed to various anthelmintic compounds (10 μM) or vehicle control. Each sheet corresponds to a specific compound and includes parameters such as raw fluorescence intensity, calculated metrics (e.g., r and K), ratio to control, and number of animals analyzed. The dataset also includes results of statistical analyses (t-tests), thresholds for toxicity classification, and annotations regarding compound toxicity (e.g., TOXIC, NEGATIVE, ACUTE). This comprehensive dataset supports the characterization of drug-induced developmental and physiological perturbations *in vivo*.

**S3 Table: Summary of data collected from the in vitro larval development and migration trap assays using N2 and SJ4100 *C. elegans* strains.** This dataset presents raw and normalized values obtained from two phenotypic assays evaluating developmental and neuromuscular function in *C. elegans* wild-type (N2) and SJ4100 strains. The LDA quantifies developmental progression based on motility after 48 hours of exposure to serial dilutions of eight different anthelmintics, reporting raw readouts, calculated efficacy percentages, and median values per condition. The MTA evaluates locomotor behavior after 24 hours of drug exposure by measuring the worms’ ability to migrate, with efficacy calculated as the percentage reduction in migration compared to controls. Each assay was independently performed on both strains, enabling the comparison of drug-induced effects on development and motility across genetic backgrounds.

## Author Contributions

CB, MB and AV performed the experiments. LR, TBD, and LM conceived of and planned the experiments. TBD wrote the manuscript with the support of LR, MB, AV, and LM. All authors provided critical feedback and helped shape the research and data analysis.

